# Downregulation is the dominant effect of new regulatory mutations in a fungal pathogen

**DOI:** 10.64898/2026.03.06.710201

**Authors:** Ana Margarida Sampaio, Daniel Croll

**Affiliations:** Laboratory of Evolutionary Genetics, Institute of Biology, University of Neuchâtel, 2000 Neuchâtel, Switzerland

## Abstract

Well-tuned gene regulation is essential for an organism’s survival. However, the origin and predominant effects of regulatory mutations remain poorly understood outside of model organisms. Here, we analyzed a large panel of genome sequencing data of the major fungal wheat pathogen *Zymoseptoria tritici* to recapitulate the evolutionary history of *cis*-regulatory mutations. We found that new mutations predominantly cause downregulation of the associated genes. The dominance of downregulation is reinforced for mutations occurring closest to the coding sequence and downstream. Mutations causing the strongest downregulation segregate at high frequencies in populations despite their recent origin. This suggests that selection may have played a role in their rapid increase since speciation. Overall, our study highlights the power of mapping populations combined with genomic surveys to unravel fundamental patterns of regulatory evolution.

## Introduction

The regulation of gene expression is essential for the development and survival of organisms. Although organisms share conserved mechanisms of gene expression regulation, variation in gene expression profiles within and between species is common [1]. Genomic regions identified as expression quantitative trait loci (eQTLs) are important determinants of mRNA level variation within species [2–6]. eQTLs enable the association of DNA sequence variation with expression variation and, ultimately, the expression of complex traits. Most eQTLs act in *cis*, regulating the expression of neighboring genes, typically resulting in larger gene expression effect sizes than *trans*-eQTLs, which affect distant loci across the genome [5,7]. The high prevalence of *cis*-eQTLs segregating within species and their typically strong effect highlight their importance in regulatory evolution over short timespans. However, the origin and effects of mutations underpinning eQTLs within a species have largely remained unexplored.

Across the mutation spectrum, deletions and loss-of-function are generally more frequently observed in contrast to insertions or gene duplications with dosage effects [8–10]. This is at least in part explained by the likelihood to acquire such mutations, however tolerance of the organism to the mutational effects is also important. New regulatory mutations may also more likely disrupt transcriptional activation rather than causing a *de novo* gain in transcription, which leads to an overrepresentation of down-regulation effects [11–13]. Furthermore, large-effect mutations on transcription may also on average be deleterious and selection should maintain such mutations at low frequencies by purifying selection [14,15]. However, compensatory evolution may stabilize new *cis*-regulatory mutations by favoring *trans* mutations to reduce deleterious effects [16,17].

Rapid regulatory evolution was shown to occur in filamentous fungal pathogens such as the wheat pathogen *Zymoseptoria tritici*. To respond to strong environmental pressures, including the need to cope with host resistance and fungicide applications, plant pathogens have evolved a large body of regulatory mutations associated with gene expression variation. High genetic diversity and regulatory variation, in particular, is a hallmark of *Z. tritici* from the individual field scale to global diversity patterns [18–21]. A large body of *cis*-eQTLs were identified using a mapping population of field-collected isolates. Nearly two-thirds of all genes were found to segregate regulatory variants [22]. Most eQTLs were found within 2 kb up- and downstream of transcription starting sites (TSS) highlighting a dense network of regulatory mutations. The origin of the regulatory mutations underpinning the eQTLs remains unexplored though.

Here, we characterized how newly arising *cis*-regulatory mutations impact gene expression at the population scale. Specifically, we assessed whether new mutations exhibit a predominant direction of expression change (down *vs*. upregulation) and evaluated their genomic distribution across gene categories and distances to associated genes.

## Results

We assessed the evolutionary history of regulatory mutations mapped in a field population of *Z. tritici*. Overall, *cis*-eQTLs were identified for 65.3% if all genes [22]. Here, we investigated the origin and distribution of *cis*-regulatory mutations in a global collection of *Z. tritici* genomes covering all major regions where the pathogen has been identified [18]. We assessed the ancestral state at all *cis*-eQTL loci using genotypic information from the closest known sister species *Zymoseptoria pseudotritici* (Supplementary Table S2). We then examined whether the derived allele (*i*.*e*. new mutation) was associated with up- or downregulation of the nearby gene in the *cis*-eQTL mapping population. Using a large global sequencing panel, we quantified patterns of regulatory effects of the derived alleles at loci previously identified as *cis*-eQTLs (Figure 1).

**Figure 1:**
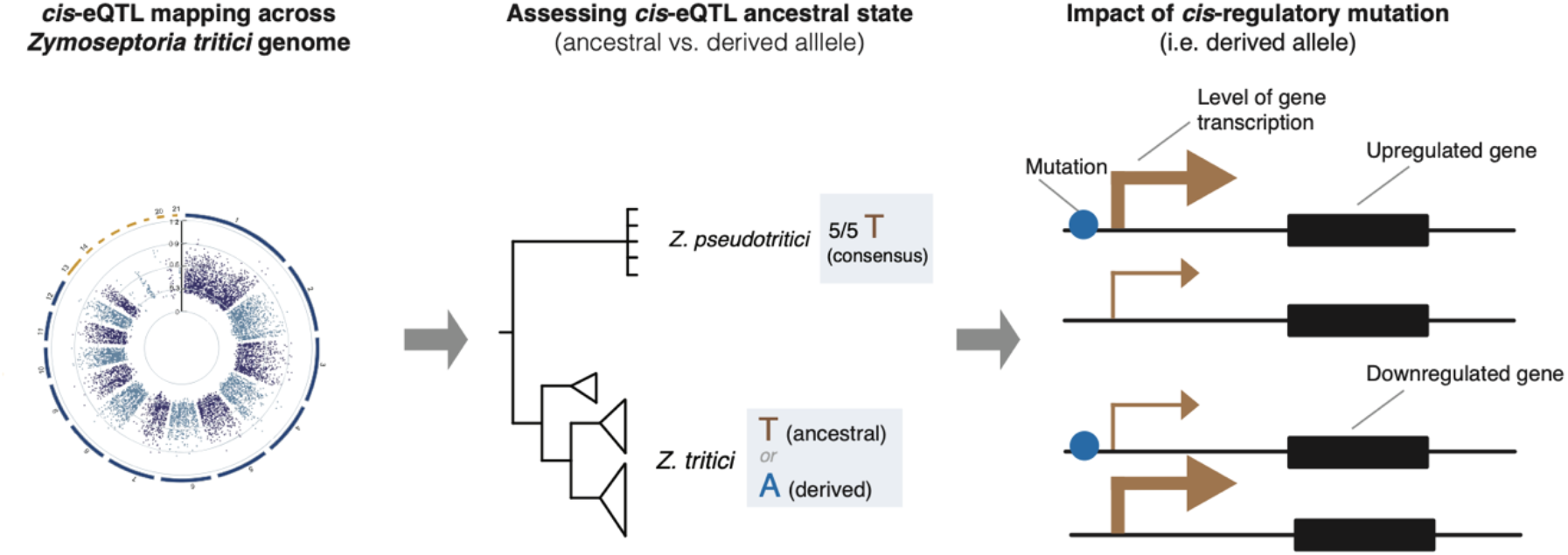
Schematic overview of the study design. *cis*-eQTLs have been mapped previously in a population of 146 *Zymoseptoria tritici* strains collected from an infected wheat field in Switzerland [22]. The circos plot identifies each significantly associated SNP locus with neighboring gene expression (*i*.*e*. eQTLs). *cis*-eQTL ancestral states were assessed based on consensus genotypes in the sister species *Z. pseudotritici*. The impact of new mutations (*i*.*e*. derived alleles) in the associated genes was examined in a large genome panel.

Derived alleles were more frequently associated with downregulation of the associated gene in the *cis*-eQTL mapping population. This pattern was even stronger in the global collection of genomes, where 82% of the new mutations (*i*.*e*. derived alleles) were associated with downregulation (Figure 2A). To evaluate whether downregulation was a dominant feature of new mutations, we examined expression associations across gene categories and the distance between the eQTL and the target gene in the global collection of genomes (Supplementary Table S3). Downregulation was similarly common for genes on core (82%) and accessory (85%) chromosomes (Figure 2B). Similarly, downregulation was the most frequent effect of new mutations across different associated gene categories (Figure 2C). Interestingly, secondary metabolite gene clusters showed the most discrepancies to the genome-wide trend with new mutations being about twice as likely to cause upregulation compared to the genome-wide average (Figure 2C). New mutations located downstream of the TSS gene exhibited more frequent downregulation effects compared to mutations located upstream of the TSS (Figure 2D). Furthermore, downregulation effects of new mutations were most dominant within 2 kb downstream of the TSS (Figure 2E; Supplementary Figure S1). Upregulation effects of new mutations increased both upstream and downstream of this interval. Hence, beyond this interval harboring most *cis*-eQTLs [22], mutations in this interval also most frequently lead to downregulation.

**Figure 2.**
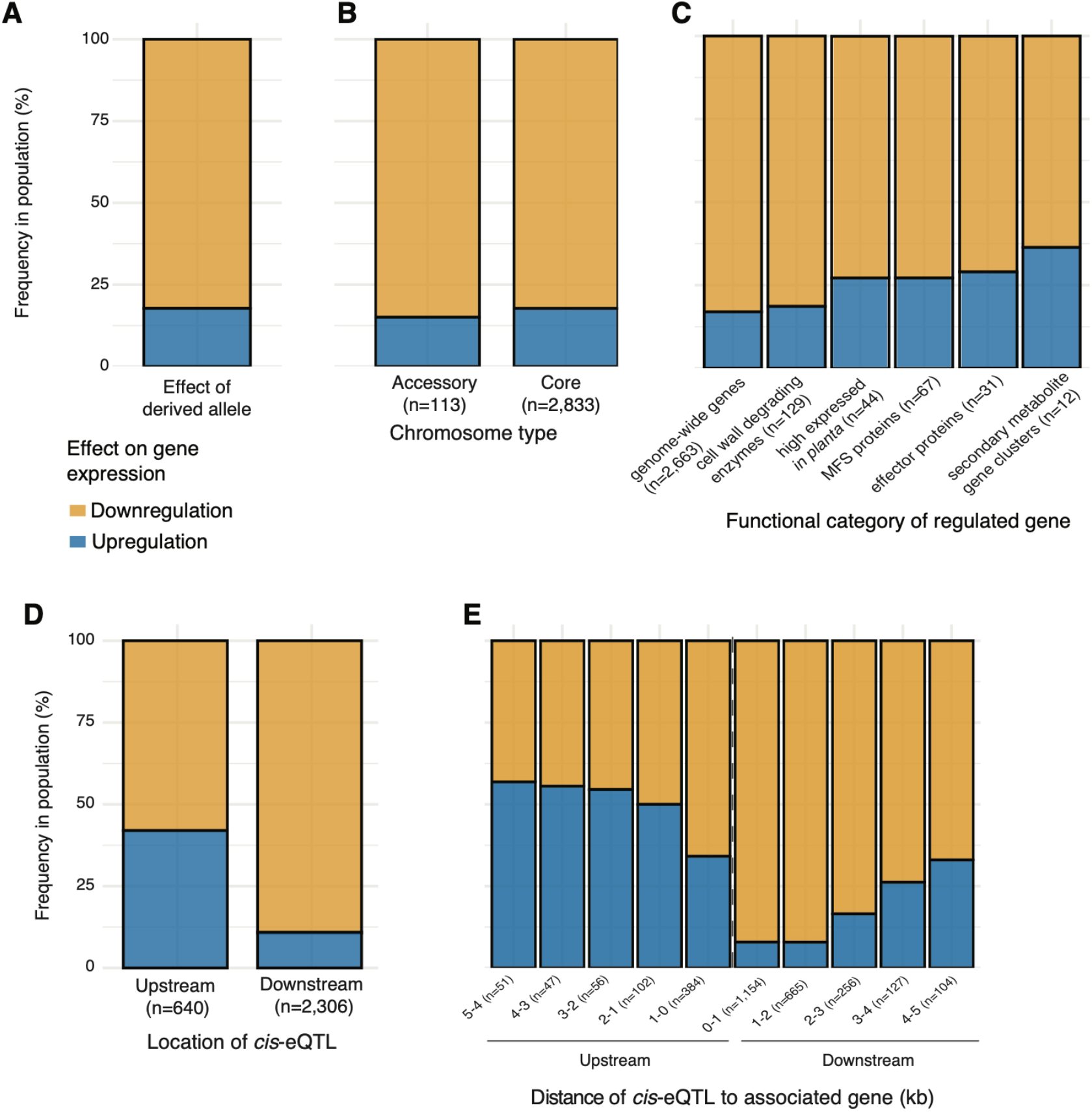
Frequency of down *vs*. upregulatory effects of new mutations at loci previously identified as *cis*-eQTL loci in the collection of genomes. A) Overall effect of derived alleles on gene expression and separated by B) chromosome type (core versus accessory), C) regulated gene functions, D) up-versus downstream position of the regulatory mutation, E) specific locations of the regulatory mutation in 1-kb windows up- and downstream of the associated gene. *n* indicates the number of derived alleles per category.

Mutations with large effects on gene expression tend to occur at lower frequencies than expected under neutral evolutionary models [14,15]. Accordingly, we expected derived alleles causing strong downregulation to be rare. To address this question, we assessed how expression effects and allele frequencies correlate in the *cis*-eQTL mapping population (Supplementary Table S4). New mutations causing the strongest downregulation were at the highest frequencies in the mapping population (Figure 3). Hence, mutations causing the strongest downregulation have reached the highest frequencies, suggesting that they are more likely to become fixed. This creates a scenario where eQTLs with the strongest effects on expression show low minor allele frequencies (Figure 3; Supplementary Figure S2). We expanded these analyses to the global collection of isolates. For this, we selected *cis*-eQTL variants with the strongest downregulation effects (effect size < -0.8). We found a similar pattern to what was observed in the Swiss population, with *cis*-eQTLs exhibiting stronger regulatory effect sizes have a derived allele at high frequency in most clusters (Supplementary Figure S3). Interestingly, one of these *cis*-eQTL loci (1_4639299) has been previously reported as contributing to local adaptation [19]. The distribution of derived alleles at this locus suggests that the mutation was beneficial in North-America and Oceania, which were colonized after the pathogen’s expansion from the Middle East to Europe [18].

**Figure 3.**
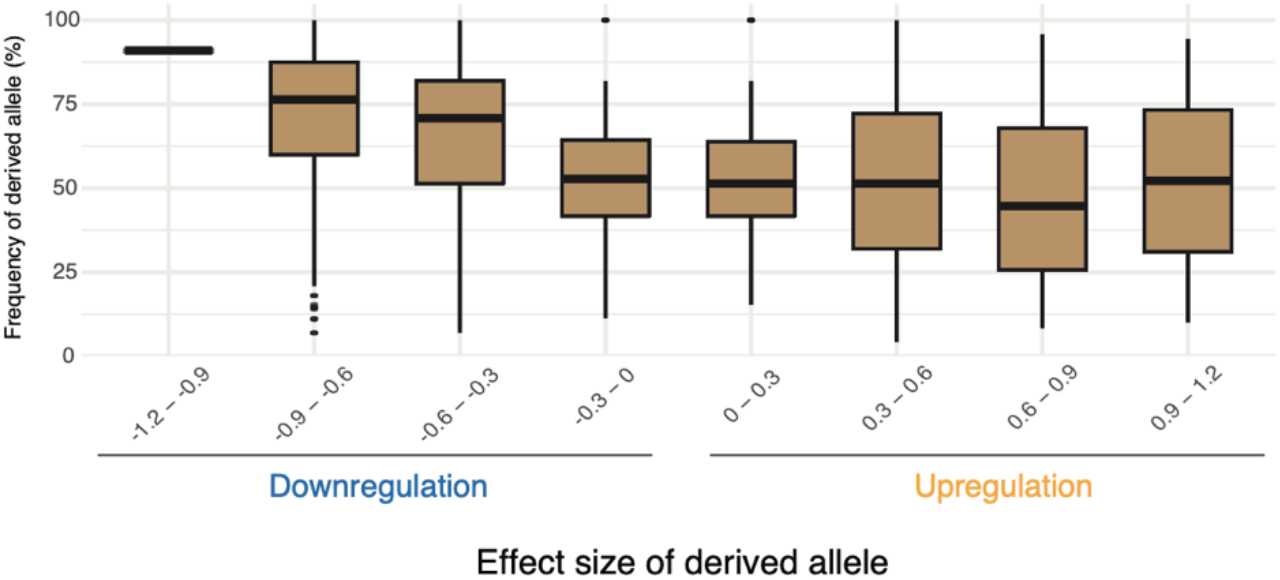
Effect sizes on gene expression of eQTL loci in the Swiss *cis*-eQTL mapping population. A) Relationship between gene expression effect size and derived allele frequency, and B) and minor allele frequency.

## Discussion

Here, we characterized how newly arising *cis*-regulatory mutations impact gene expression in a fungal pathogen species. Our analyses show that new mutations predominantly cause downregulation of the associated genes. This effect is exacerbated if the regulatory mutation occurred downstream and in close proximity to the coding sequence. Interestingly, new mutations causing strong downregulation are at high frequency in the population. Hence, mutations causing downregulation might have been favored by selection to reach high frequencies.

The observed bias toward downregulation is consistent with the fact that transcriptional disruption is more readily achieved by mutation than transcriptional enhancement [11–13]. Mutations with large effects are expected to be rare [14,15]. Consistent with this expectation *cis*-eQTLs of large effects tend to have low allele frequencies in the mapping population [22]. However, selection might act more strongly against expression increases, since they can impose higher metabolic cost or causes toxicity, reducing organism fitness [23,24], whereas moderate down-regulation may be more tolerated. Our analyses of ancestral states at eQTLs revealed that down-regulation was indeed the dominant effect of new mutations (*i*.*e*. derived alleles). Furthermore, these mutations rose to high frequency in the mapping population, as well as broadly across the pathogen’s global distribution range. Under stabilizing selection, large deviations from optimal expression, either up or down, should generally be deleterious and therefore rare. Hence, new mutations with strong effects should have remained at low frequencies rather than gaining such high frequencies. This suggests that the strong downregulation mutations may represent variants of weak or no fitness consequences. A small subset may have been favored by selection if strong down-regulation was beneficial.

We analyzed positional effects of *cis*-regulatory mutations causing downregulation and found that these rose to higher population frequency if the eQTL was located downstream and close (< 2 kb) to the associated gene. These gene proximate regions are strongly enriched for *cis*-eQTLs [22]. More distal regulatory variants may more often be buffered by chromatic architecture or other epigenetic mechanisms [25,26]. The population frequency of down-regulation mutations was also associated with the function of the regulated genes. Genes encoding secondary metabolite clusters and candidate effectors involved in host interactions, were more likely to have gained a mutation causing upregulation compared to other categories, where downregulation is the dominant effect of new mutations. This observation may be consistent with several possible explanations. First, secondary gene clusters and pathogenicity-associated genes tend to be more likely controlled by epigenetic modifications rather than by changes in transcription factor binding, for example [27,28]. Such differences could have an impact on the likelihood of a new mutation to upregulate gene expression. Mutations could be associated, for example, with the insertion or excision of a transposable element with effects on chromatin organization [29]. Second, the emergence of *Z. tritici* as a major wheat pathogen from an ancestor with a possibly more endophytic lifestyle [30]. This transition could have selected for the upregulation of secondary gene clusters and other pathogenicity-related genes. Molecular genetics analyses will be required to disentangle exact roles of regulatory mutations on genes and their causal role in the pathogen lifestyle.

In conclusion, we show that new *cis*-regulatory mutations predominantly result in gene downregulation and that such mutations have risen to high frequency. This highlights the power of combined population genomics and eQTL mapping approaches to investigate broad patterns of regulatory evolution.

## Material and Methods

### Expression QTL mapping population

A genome-wide map of regulatory polymorphisms governing gene expression has been established based on a collection of 146 *Z. tritici* isolates collected from a single field site in Switzerland [22]. Briefly, trimmed high-quality DNA short read sequences were aligned to *Z. tritici* reference genome IPO323 [31] using Bowtie2 (v2.3.4.3) [32] and used for variant calling based on the HaplotypeCaller integrated in GATK (v4.0.11.0) [33]. SNPs were filtered for quality with indels and multialelic sites being removed using bcftools (v1.9) [34]. Detailed filtering steps were reported previously [22]. RNA-seq datasets were trimmed and aligned to the *Z. tritici* reference genome IPO323 [31] using HISAT2 (v2.1.0) [35] with the parameter “– RNA-strandedness reverse”. By combining SNP calling data with expression data, *cis*-eQTLs were mapped using QTLtools (v 1.1) [36] using the *cis* conditional option and choosing a *cis* window of 10 kb equidistant from the TSS. SNPs associated with mRNA abundance of a single, proximate gene were defined as *cis* eQTLs and used for further analyses.

### Global thousand-genome resequencing panel

We assessed global frequencies of regulatory mutations based on a large population sequencing panel covering all major wheat-producing regions where the pathogen is present. The thousand-genome panel comprises 1035 genomes (Supplementary Table S1) [18]. Variant calling was performed as described earlier [18] with SNPs mapped to the IPO323 reference genome using Bowtie2 (v2.4.1) [32]. We classified the associated *cis*-eQTL alleles as ancestral or derived using genome sequencing data of the closely related sister species *Z. pseudotritici*. For this, we used publicly available PacBio long-read sequencing data from five *Z. pseudotritici* isolates from the species endemic region in Iran (NCBI SRA accessions SRR11972499-SRR11972503). Reads were aligned to the reference genome IPO323 [37] using minimap2 (v2.30) [38]. Variant calling was performed using bcftools (v1.22) [34]. Ancestral and derived allele states were assigned only at SNPs showing a consistent allele state across all five *Z. pseudotritici* isolates. The *Z. pseudotritici* allele was considered to be ancestral. *cis*-eQTL transcriptional effect sizes were obtained from QTLtools. Minor alleles were determined based on allele frequencies in the thousand-genome global panel.

## Author contributions

AMS and DC conceived the study. AMS performed analyses. AMS and DC wrote and revised the manuscript.

## Data availability

All sequence data is deposited on the NCBI Short Read Archive under the BioProject accession number PRJNA650267 (https://www.ncbi.nlm.nih.gov/bioproject/PRJNA650267).

## Acknowledgments

We are grateful to Leen Nanchira Abraham for assistance and helpful advice. The study was supported by Swiss National Science Foundation grant to DC (201149).

## Notes

### Competing Interest Statement

The authors have declared no competing interest.

