## Supplementary Figures for "Downregulation is the dominant effect of new regulatory mutations in a fungal pathogen"

Sampaio et al.

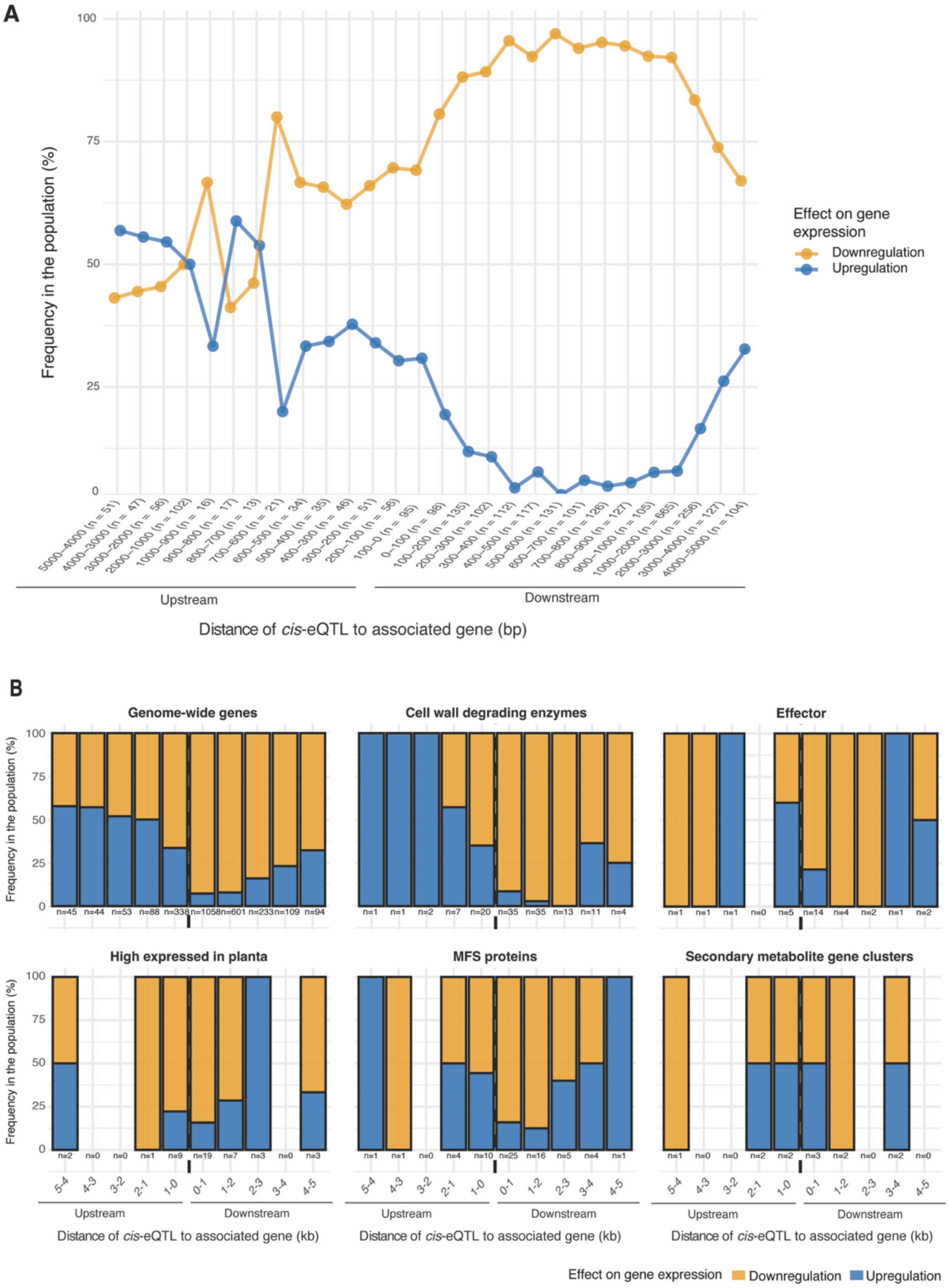


**Supplementary Figure S1**: Frequency of down *vs.* upregulation effects of new mutations at loci previously identified as *cis*-eQTL loci in the global genome panel by: A) *cis*-eQTL distance to associated gene. B) *cis*-eQTL distance to associated gene, per regulated gene functions.

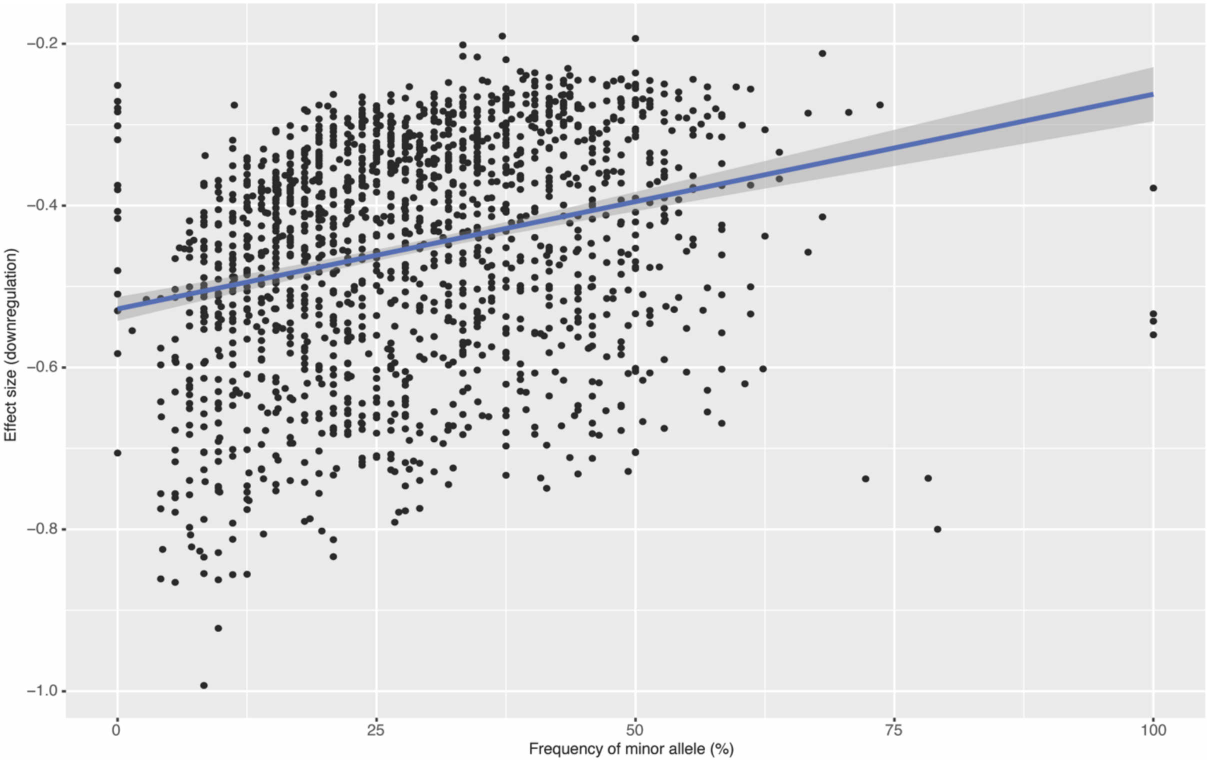


**Supplementary Figure S2**: Correlation between downregulation effect size and minor allele frequency in the Swiss population.

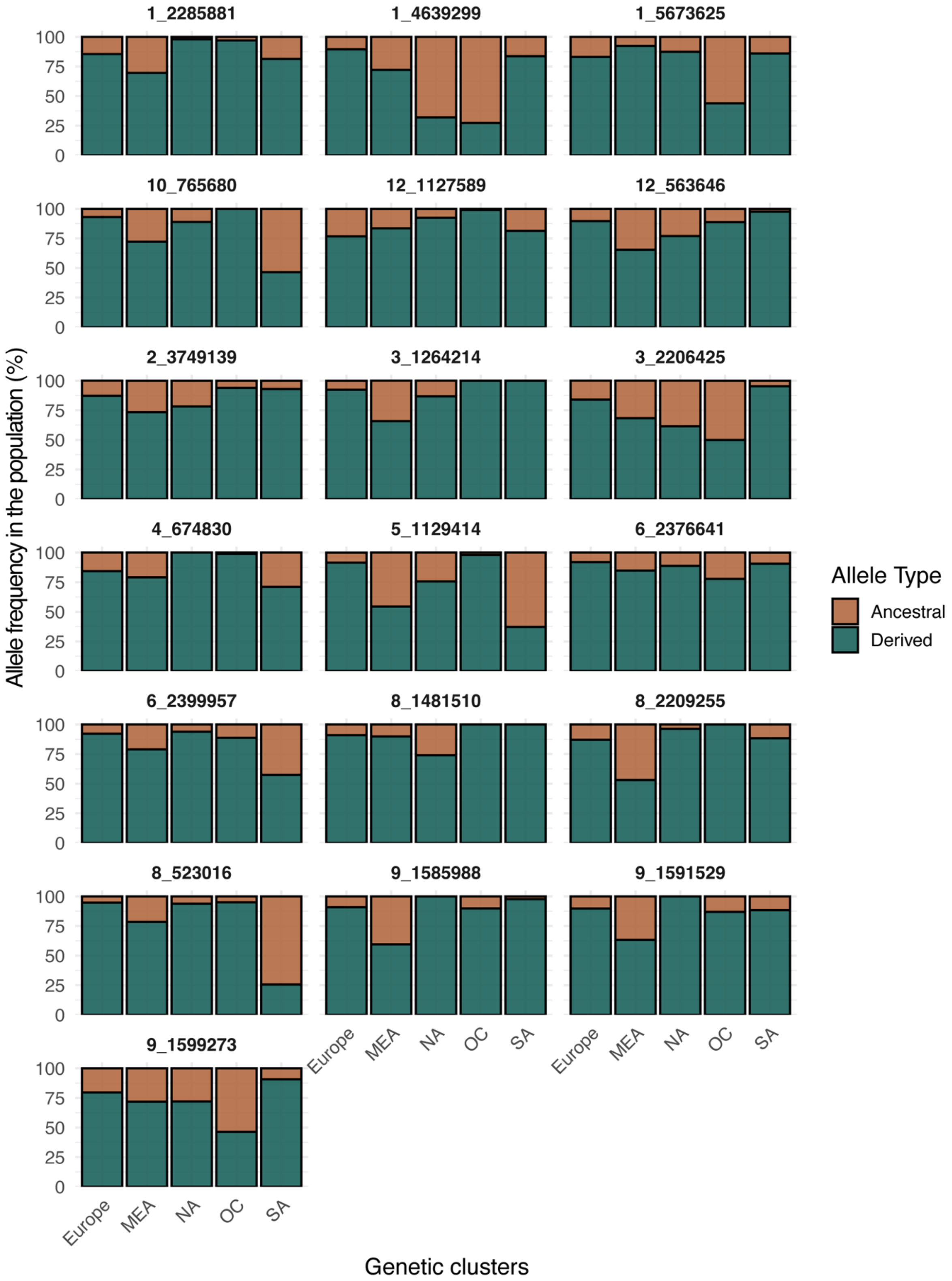


**Supplementary Figure S3**: Frequency of ancestral and derived alleles at *cis*-eQTLs with stronger down-regulation effect sizes. Numbers indicate chromosome and respective position of the alleles. Genetic clusters: MEA – Middle East, NA – North America, OC – Oceania, SA – South America.
